# EASTR: Correcting systematic alignment errors in multi-exon genes

**DOI:** 10.1101/2023.05.10.540179

**Authors:** Ida Shinder, Richard Hu, Hyun Joo Ji, Kuan-Hao Chao, Mihaela Pertea

## Abstract

Accurate alignment of transcribed RNA to reference genomes is a critical step in the analysis of gene expression, which in turn has broad applications in biomedical research and in the basic sciences. We have discovered that widely used splice-aware aligners, such as STAR and HISAT2, can introduce erroneous spliced alignments between repeated sequences, leading to the inclusion of falsely spliced transcripts in RNA-seq experiments. In some cases, the “phantom” introns resulting from these errors have made their way into widely-used genome annotation databases. To address this issue, we have developed EASTR (Emending Alignments of Spliced Transcript Reads), a novel software tool that can detect and remove falsely spliced alignments or transcripts from alignment and annotation files. EASTR improves the accuracy of spliced alignments across diverse species, including human, maize, and *Arabidopsis thaliana*, by detecting sequence similarity between intron-flanking regions. We demonstrate that applying EASTR before transcript assembly substantially reduces false positive introns, exons, and transcripts, improving the overall accuracy of assembled transcripts. Additionally, we show that EASTR’s application to reference annotation databases can detect and correct likely cases of mis-annotated transcripts.

## Introduction

RNA sequencing (RNA-seq) is a widely used method for quantifying gene expression and characterizing transcriptome diversity. However, repetitive sequences can in some circumstances induce splice-aware aligners, such as STAR and HISAT2, to create spurious introns spanning two nearby repeats. Repeat elements constitute a significant portion of many genomes, comprising 21% of the *Arabidopsis thaliana* genome [1], 53% of the human genome [2], and 85% of the *Zea mays* genome [3]. Repeated elements frequently occur in close proximity; for example, *Alu* elements, a primate-specific transposable element (TE), appear in over a million copies in the human genome, with an average frequency of once every 3,000 bases [4]. The close proximity of repeat elements complicates distinguishing spliced and contiguous alignments, particularly in tissues and organisms with high TE expression.

Repeat elements pose challenges not only due to their proximity but also due to their high degree of polymorphism. The variations among individuals and between loci can confound computational methods attempting to distinguish correct and incorrect spliced alignments. As we will show below, this can result in the inclusion of transcripts with spurious junctions in human gene catalogs such as CHESS [5], RefSeq [6], and GENCODE [7].

Computational methods face inherent limitations when aligning short reads from sequencing data to genomes with numerous repeat elements. Read lengths are often shorter than the full length of repeated sequences, complicating the identification of the true origin of multimapped reads. Pseudogenes also present challenges during alignment, as reads that should be aligned across a splice site at their original location may be aligned end-to-end to a pseudogene copy. HISAT2 [8] addresses this by prioritizing spliced alignments over contiguous alignments when mapping quality is similar (**Figure 1**). However, this approach can lead to misalignments of other types of repeats such as transcripts with variable numbers of tandem repeats (VNTRs) or TEs that have accumulated variation and diverged in sequence over evolutionary time.

**Figure 1.**
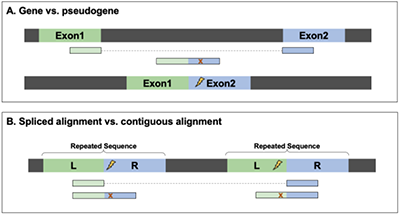
Assessing HISAT2 algorithm’s performance in two alignment scenarios: **(A)** The correct spliced alignment without mismatches at a gene locus is favored over an unspliced alignment with one mismatch to a pseudogene. HISAT2 accurately aligns a read (blue and green rectangles) originating from a gene with exon 1 (green) and exon 2 (blue), which also aligns contiguously to a processed pseudogene on a different chromosome. The read contains a single mismatch (marked with an x) to the pseudogene. When the read is aligned to its correct location, spanning an intron, the alignment has zero mismatches. **(B)** An incorrect spliced alignment caused by consecutive repeats can occur when a read (blue and green rectangles) originates from either repeat 1 or repeat 2, but due to either a basecalling error or a polymorphism, it contains mismatches (marked with x) to both repeat elements. The correct alignment is end-to-end, spanning an entire repeat. Because the repeat’s left arm (marked L) ends with AG and the right arm starts with GT (introducing a sequence resembling a canonical splice site), the aligner can generate a false intron and align the read without mismatches.

To address the issue of incorrect spliced alignment between repeated sequences, we developed EASTR (Emending Alignments of Spliced Transcript Reads), a tool that detects and removes erroneous spliced alignments by examining the sequence similarity between the flanking upstream and downstream regions of an intron and the frequency of sequence occurrence in the reference genome.

## Results

To demonstrate the versatility and applicability of EASTR, we applied it to three model organisms with distinct genomic repeat content and to samples with different library preparation methods. The organisms and tissues we selected were human brain [9], *Zea mays* (maize) leaves [10] and pollen [11], and *Arabidopsis thaliana* [12]. Our objective was to highlight the effectiveness of EASTR in enhancing alignment accuracy across a broad range of organisms and experimental designs.

To evaluate the impact of EASTR alignment filtering on downstream analyses, we assessed the accuracy of transcript assembly before and after filtering the alignments. We used StringTie2 [13] to assemble transcripts from both HISAT2 and STAR [14] alignments, as well as from the EASTR-filtered alignments. Our results described below demonstrate that filtering with EASTR prior to assembly improved both sensitivity and precision, and reduced the number of non-reference introns, exons, and transcripts, which are more likely to represent transcriptional noise [15].

In addition to filtering alignment files, we also used EASTR to identify potentially erroneous transcripts in reference annotation databases for each of the selected model organisms. By examining the sequence similarity between the flanking upstream and downstream regions of introns, EASTR was able to detect transcripts in the annotation that may have been incorrectly annotated due to spliced alignment errors between repeat elements.

### Human

We evaluated EASTR’s performance on paired RNA-seq datasets from developing and mature human dorsolateral prefrontal cortex (DLPFC). The datasets were obtained using both poly(A) selection and rRNA-depletion (ribo-minus) library preparation methods from cytoplasmic and nuclear fractions from three prenatal and three adult samples [9].

#### Reducing Spurious Junctions in Alignments

We applied EASTR to identify putative erroneous junctions in the alignment files of 23 DLPFC samples. Our analysis revealed that the vast majority of the alignments flagged for removal by EASTR supported junctions not present in the RefSeq reference annotation. On average, EASTR removed 3.4% (5,208,893/153,192,435) and 2.7% (3,599,371/134,202,142) of all HISAT2 and STAR spliced alignments, respectively. Of the removed alignments, only 0.2% (9,114) in HISAT2 and 0.3% (9,101) in STAR supported 114 and 119 reference-matching junctions, respectively. EASTR marked these as erroneous in the RefSeq reference annotation, and this small subset is discussed further in the section on the application of EASTR to reference annotations. Nearly all of the alignments targeted for removal by EASTR were at non-reference junctions: 99.8% (5,199,779) in HISAT2 and 99.7% (3,590,270) in STAR, corresponding to 138,111 and 75,273 non-reference junctions, respectively. This reduction in the number of non-reference junctions was consistent across all 23 samples. More details are provided in **Supplemental Table S1.1** and **Figure 2**.

**Figure 2.**
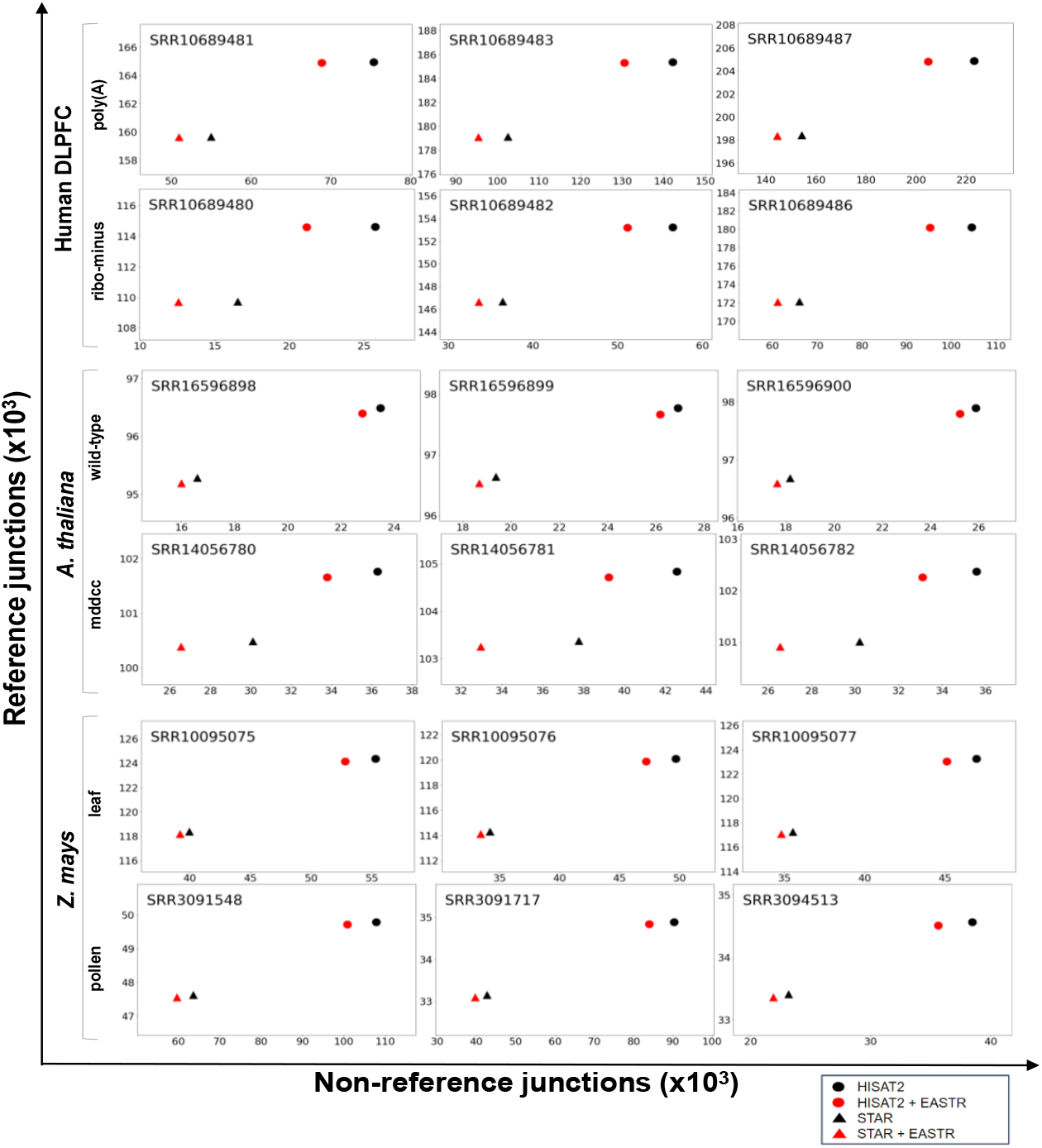
Comparison of reference and non-reference junction counts before and after EASTR filtering, illustrating the effectiveness of EASTR filtering in distinguishing between reference and non-reference junctions across various sample types. We chose three samples from each dataset, including human DLPFC polyA and ribo-minus, *A. thaliana* wild-type and *mddcc* strains, and *Z. mays* lower leaf and mature pollen. The y-axis indicates the count of junctions matching the reference annotation for a given sample, whereas the x-axis shows the count of junctions not present in the reference annotation.

We found that the ribo-minus library method had a higher proportion of spuriously spliced alignments compared to the poly(A) selection method. Of the 23 samples, 11 pairs were processed using both library selection methods. In ribo-minus samples, EASTR flagged 8.0% (4,145,349/51,742,668) and 6.4% (2,481,034/39,030,763) of HISAT2 and STAR alignments as erroneous, respectively, compared to only 1.0% (1,063,544/101,449,767) and 1.2% (1,118,337/95,171,379) in poly(A) samples. Furthermore, our findings suggest that developmental stage may be a relevant factor to consider during alignment and downstream RNA-seq analysis. Comparing ribo-minus adult to ribo-minus neonatal samples revealed that, in general, prenatal samples had a higher rate of removed spliced alignments in comparison to adult samples (**Supplemental Table S1.1**).

#### Improving Transcript Assembly Quality

We assembled transcripts for each sample using StringTie2 from HISAT2 and STAR alignments, both unfiltered and filtered by EASTR. We compared the resulting transcript assemblies to the RefSeq human reference annotation and found that filtering with EASTR improved transcript assembly quality, as summarized in **Table 1**. The decrease in the number of non-reference introns and exons, as well as the relative improvement in transcript-level precision, can be observed across all samples and alignment tools (**Supplemental Tables S2.1** and **S3.1**). Importantly, aligning with HISAT2 and subsequently filtering with EASTR did not compromise transcript-level sensitivity in any sample or experimental condition and even resulted in slight improvements relative to the reference annotation. When aligning with STAR and then filtering with EASTR, the decline in transcript-level sensitivity was almost zero, affecting only 1-2 transcripts in two samples. Our results strongly support the adoption of EASTR to enhance transcriptome assembly precision while preserving or improving sensitivity.

**Table 1.** Summary of the impact of EASTR filtering on transcriptome assembly metrics for different species and conditions, comparing assemblies generated from unfiltered alignments to those filtered with EASTR. The table shows the percentage change (%Δ) in the number of non-reference introns and non-reference exons, and the change in count (Δ) of non-reference and reference transcripts.

| A. HISAT2 | | Non-reference introns %$\Delta$ | Non-reference exons %$\Delta$ | Non-reference transcripts $\Delta$ | Reference transcripts $\Delta$ |
| --- | --- | --- | --- | --- | --- |
| Human DLPFC | ribo-minus | -5.1% to -22.1% | -3.7% to -19.8% | -148 to -1,076 | +0 to +52 |
|  | polyA | -6.1% to -9.1% | -4.3% to -7.7% | -199 to -797 | +3 to +135 |
| A. thaliana | mddcc | -3.6% to -7.4% | -7.1% to -8.2% | -119 to -176 | -7 to -11 |
|  | wild-type | -5.0% to -5.8% | -7.2% to -9.3% | -86 to -94 | -8 to -14 |
| Z. mays | leaf | -2.1% to -3.5% | -2.5% to -3.2% | -232 to -264 | -23 to -43 |
|  | pollen | -3.5% to -6.2% | -3.8% to -5.4% | -237 to -456 | +5 to +12 |

| B. STAR | | Non-reference introns %$\Delta$ | Non-reference exons %$\Delta$ | Non-reference transcripts $\Delta$ | Reference transcripts $\Delta$ |
| --- | --- | --- | --- | --- | --- |
| Human DLPFC | ribo-minus | -3.7% to -14.3% | -2.3% to -12.9% | -89 to -603 | -2 to +42 |
|  | polyA | -5.3% to -7.7% | -3.2% to -6.0% | -162 to -561 | +7 to +86 |
| A. thaliana | mddcc | -2.7% to -3.2% | -5.6% to -6.0% | -87 to -100 | -5 to -13 |
|  | wild-type | -4.9% to -6.3% | -8.1% to -8.9% | -56 to -68 | -11 to -19 |
| Z. mays | leaf | -1.0% to -1.3% | -1.2% to -1.8% | -135 to -185 | -32 to -35 |
|  | pollen | -1.2% to -2.1% | -1.3% to -1.9% | -65 to -140 | +2 to +10 |

#### Evaluating reference annotation accuracy

We also used EASTR to assess the accuracy of junctions in widely used human reference annotation catalogs, including RefSeq (version 110), CHESS (version 3.0), GENCODE (version 41), and MANE (version 1.0) [16]. EASTR detected 365 potentially spurious introns across 581 transcripts and 237 genes in RefSeq, 192 introns across 319 transcripts and 124 genes in CHESS, and 411 introns across 475 transcripts and 344 genes in GENCODE (**Supplementary Table S4.1-S4.3**). Notably, we also identified one incorrect MANE transcript, as discussed below.

Our investigation highlighted that gene families characterized by frequent gene duplication, complex repetitive structures, and variable copy number across individuals and populations are particularly susceptible to splicing errors in reference annotations. For instance, the primate-specific gene family NBPF features a two-exon repeat unit known as Olduvai that has expanded through tandem duplication [17, 18]. We analyzed the NBPF20 gene, one of the longest members of this family with a remarkable expansion of this repeat unit [17]. As shown in **Figure 3A**, the presence of the two-exon repeat unit in the NBPF20 gene presents challenges in accurately distinguishing between contiguous and spliced alignments. The RefSeq transcript NM_001278267 (CHESS transcript CHS.2819.2), displays sequence homology between exons 125_1 and 126_1, as well as the “intronic” region between them, suggesting they form two halves of a repeated exon. A comparison of shortened exons 125_1 and 126_1 in transcript NM_001278267 with their full-length counterparts in exons 124_2 and 126_2 in transcript NM_001397211 supports this assertion. As a result, spliced alignments supporting exons 125_1 and 126_1 also align contiguously to exons 124_2 or 126_2. Additionally, a sharp drop in coverage of spliced alignments drops abruptly at the break in sequence homology between the two paralogous exons, further indicating inaccurately spliced alignments (**Figure 3A**, coverage and alignment tracks).

**Figure 3.**
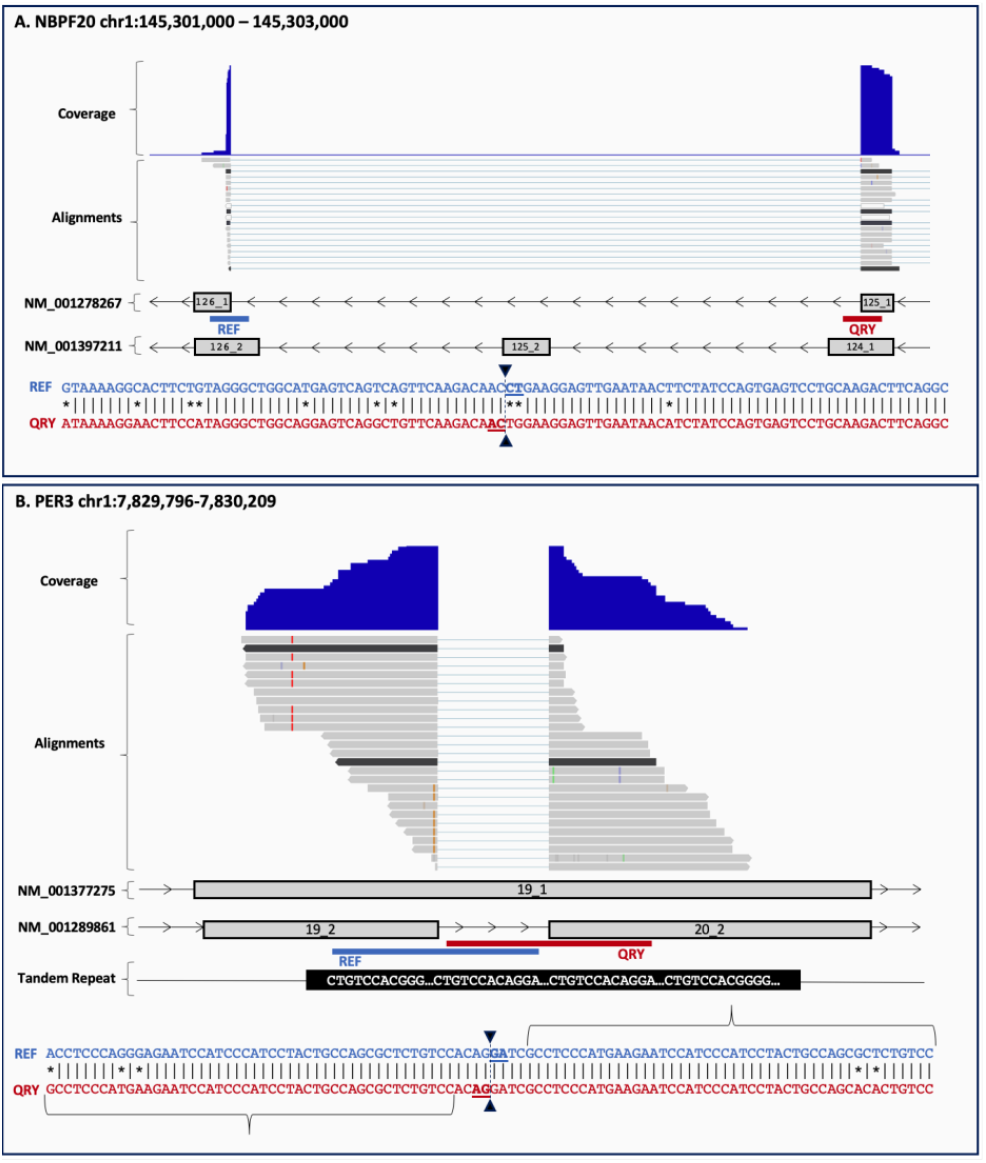
Examples of splicing errors in reference annotation transcripts caused by complex repetitive structures or polymorphisms. **Panel A** shows an error in NBPF transcript NM_001278267 where a complex repetitive structure causes an intron to be included between exons 125_1 and 126_1, skipping exon 125_2. The correct form of the transcript occurs in MANE (NM_001397211) and includes the missed exon as well as longer versions of each of the flanking exons. The repeat that causes the error includes the 100bp alignment between the upstream (REF) and downstream (QRY) flanking regions shown below the transcripts. Inverted triangles at the center of the 100bp alignment mark the splice sites. Above the transcripts in the figure, two tracks are displayed: the coverage track and the alignment track of spliced reads across 23 brain (DLPFC) samples, which were combined into a single alignment file using TieBrush [20]. In the coverage and alignment tracks, only the alignments that support the junction between exons 124_1 and 125_1 are shown. **Panel B** displays an error in PER3 transcript NM_001289861 caused by a tandem repeat. Exon 19_1 in MANE transcript NM_001377275 overlaps with a 54bp tandem repeat, and an intron of the same length is erroneously inserted between exons 19_2 and 20_2 in NM_001289861. The first 46bp of QRY and the last 46bp of REF (as shown by the braces), represent the genomic region overlap between the flanking sequences. Inverted triangles at the center of the alignment denote the splice sites. The coverage and alignment of spliced reads (alignment track) across 23 DLPFC samples further supports the presence of the error.

A substantial proportion of introns flagged as questionable by EASTR were less than 100bp in length (36% in CHESS, 45% in RefSeq, and 11% in GENCODE), coinciding with regions containing variable number of tandem repeat (VNTR) polymorphisms. For example, the PER3 gene contains a VNTR with either 4 or 5 repeated 54bp sequences [19]. In RefSeq transcript NM_001289861 (CHESS3 CHS.278.18, GENCODE ENST00000614998.4), EASTR identified a 54bp intron that matches the periodicity of the VNTR region (**Figure 3B**). Only 4 out of 27 alignments supporting this intron are without mismatches to the reference genome and have sufficient overhangs extending beyond the region of homology on both ends of the junction. These alignments are likely indicative of an indel, rather than an intron. The remaining 23 alignments exhibit either short overhangs on one end of the junction not extending beyond the homologous region or contain mismatches to the reference, or a combination of both (**Figure 3B**, alignment track).

In the MANE catalog, a recently established database that selects one isoform for each proteincoding gene to serve as the representative transcript for that gene, and on which RefSeq and GENCODE agree perfectly, we identified a TCEANC gene transcript (RefSeq: NM_001297563, CHESS: CHS.57562.1, GENCODE: ENST00000696128) containing an intron that appears erroneous (**Figure S1, Supplemental Table S4.4**). This intron features splicing between two consecutive *Alu* elements sharing 84% sequence identity, potentially causing alignment ambiguity that may be compounded by individual polymorphisms. Moreover, the inclusion of the second *Alu* element disrupts the open reading frame (ORF), shortening it substantially (**Figure S1**). The questionable intron is further characterized by a low-quality splice site acceptor motif, as depicted in **Figure S2**. Other gene catalogs exhibit many additional *Alu-Alu* splicing events: EASTR flagged 221 instances in GENCODE, 20 in CHESS, and 10 in RefSeq. Additionally, we identified an instance where the protein-coding gene NPIPB3 is absent from the MANE catalog, likely due to differences between RefSeq and GENCODE regarding the “correct” splice site within a VNTR region in final last exon (Supplemental Material, Section 2).

### Zea mays

We evaluated the performance of EASTR on six RNA-seq datasets from *Zea mays*, consisting of three biological replicates each from mature pollen and lower leaves affected by gray leaf spot disease. Previous research has shown higher TE expression in reproductive tissue compared to vegetative tissue [12], leading us to hypothesize that the mature pollen dataset would contain a larger number of spurious alignments. Our results supported this hypothesis, indicating that EASTR was more effective in identifying false alignments in the mature pollen dataset than in the lower leaf dataset.

#### Reducing Spurious Junctions in Alignments

Compared to the lower leaf dataset, EASTR identified a higher proportion of spurious alignments in the mature pollen dataset (**Figure 2**). In this dataset, EASTR flagged 12.3% (1,840,959/14,923,699) and 14.8% (1,548,226/10,467,851) of HISAT2 and STAR spliced alignments as spurious, respectively, while only 0.8% (120,215/15,177,186) and 0.4% (61,286/13,981,211) were flagged in the leaf dataset. In the pollen dataset, only a small proportion of removed HISAT2 and STAR alignments, (1.0% and 1.2%, respectively) corresponded to junctions in the reference annotation (79 for HISAT2 and 87 for STAR). The remaining 13,338 and 6,645 alignments for HISAT2 and STAR corresponded to non-reference junctions. In the leaf dataset, 8.8% and 13.6% respectively of removed HISAT2 and STAR alignments supported reference junctions, corresponding to 260 and 234 annotated splice sites, with the remaining alignments supporting non-reference junctions (5,538 and 1,786, respectively).

#### Improving Transcript Assembly Quality

Alignment filtering using EASTR had a clear positive impact on the accuracy of the subsequent transcriptome assembly with StringTie2, as shown in **Table 1**. In the pollen dataset, the number of non-reference introns, exons, and transcripts was reduced, without compromising transcript-level sensitivity or the number of reference-matching transcripts assembled (Supplemental Table S2.3 and S3.3). Filtering the leaf dataset with EASTR also resulted in a decrease in the number of non-reference introns, exons, and transcripts. While EASTR improved transcript-level precision in the leaf samples, there was a slight reduction in sensitivity (0.2-0.3% for HISAT2 and 0.2% for STAR) and in the number of reference-matching transcripts assembled (<40 removed out of >15,000 total). Taken together, these results suggest that filtering spurious alignments can result in a more precise and comparably sensitive transcriptome assembly.

#### Evaluating reference annotation

We utilized EASTR to evaluate the maize genome annotation obtained from MaizeDGB [21,22] (version 5.0 of the B73 inbred line) and identified 412 potentially spurious introns within 539 transcripts and 261 genes (**Supplementary Table S4.5**). Our analyses revealed that tandemly repeated sequences were the main cause of erroneous splice site annotation, as illustrated in **Figure 4**. In **Figure 4A** we show the annotation of a single gene with two spliced transcripts at the chr10:92,245,678-92,291,403 locus. However, analysis with EASTR suggests that these two transcripts of gene ACCO3 may represent tandemly duplicated genes rather than two transcripts of the same gene. Both transcripts are 1320 bp long and share 99.4% sequence identity. In **Figure 4B**, we illustrate a case of likely splice site mis-annotation between duplicated TEs. Although the B73 TE annotation track displays only one TE in this region, the dot plot shown in the figure indicates that the TE is repeated seven times in this region. EASTR identified four potentially spurious introns annotated within this tandem repeat region.

**Figure 4.**
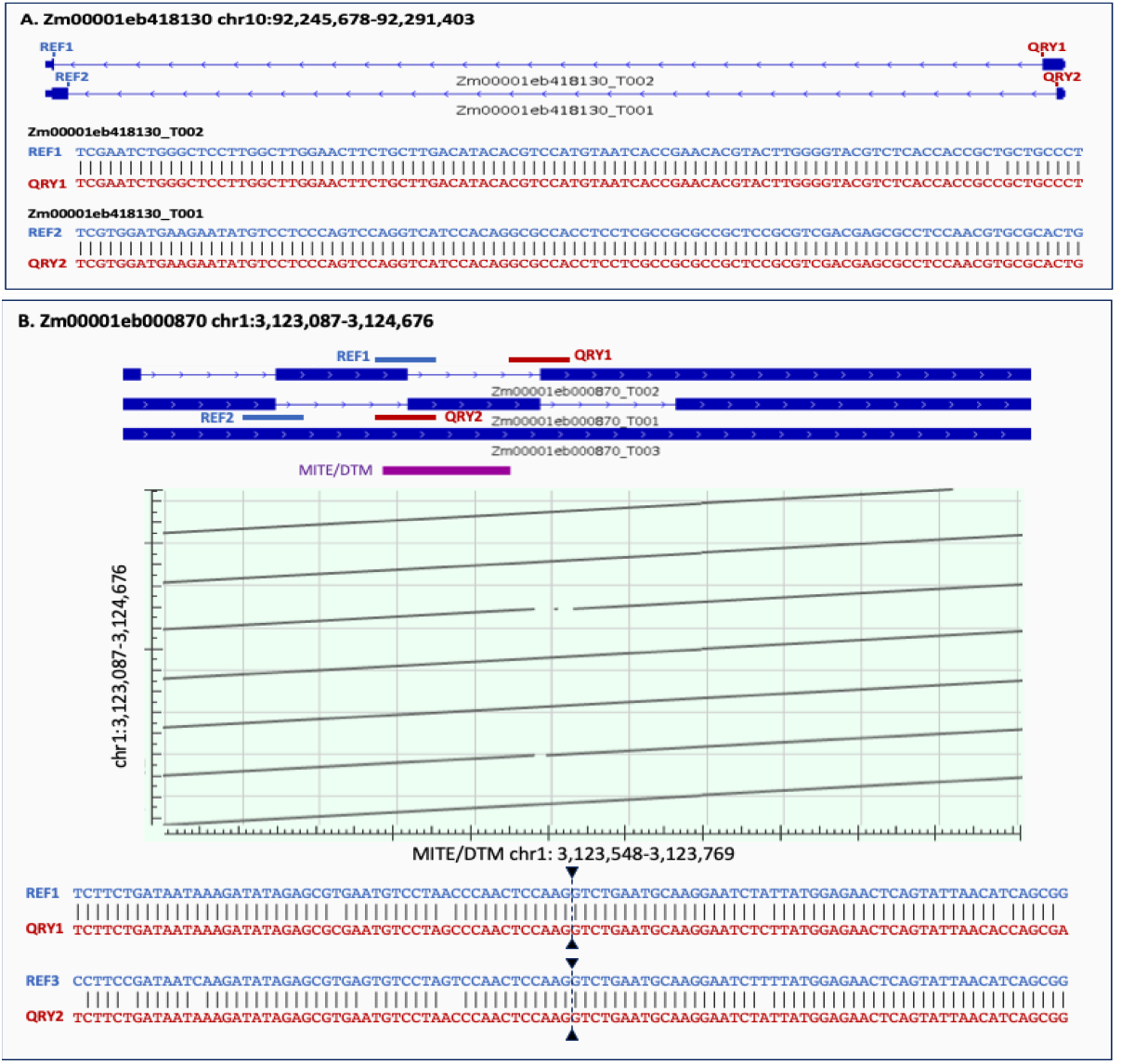
Erroneous splice site annotation between duplicated sequences: **Panel A** illustrates an instance of an error in splice site annotation involving two duplicate genes. The reference annotation (panel A, top) presents these genes as a single gene with two spliced transcripts. Alignments of the upstream and downstream intron-flanking sequences (REF1, REF2 vs QRY1, QRY2, respectively) in transcripts Zm00001eb418130_T002 and Zm00001eb418130_T001 show perfect conservation. **Panel B** presents a case of splice site annotation error between duplicated transposable elements (TEs). The top track displays three annotated transcripts, with transcript T003 being a fragment and not a full-length transcript. The TE annotation track below the transcripts shows that only a single TE is annotated in this region. However, the dot plot below the TE annotation track indicates that the annotated TE is repeated 7 times in the chr1:3,123,087-3,124,676 region. The 100 bp intron-flanking sequence alignments of two junctions in transcripts T002 and T001 are shown below the dot plot and demonstrate strong homology between the four sequences.

### A. thaliana

We evaluated EASTR’s performance on paired RNA-seq datasets from wild-type (WT) and DNA methylation-free mutant *A. thaliana* plants. The mutant plants (*mddcc*) were generated by knocking out all DNA methyltransferases (MET1, DRM1, DRM2, CMT3, and CMT2), which play an important role in maintaining DNA methylation patterns and regulating gene expression, including silencing of transposable elements (TEs) [13]. All datasets were generated using ribominus library preparation and consisted of three biological replicates for each condition. Our findings supported our hypothesis that the DNA methylation loss results in increased TE expression levels and a higher proportion of spurious spliced alignment events detected by EASTR.

#### Reducing Spurious Junctions in Alignments

EASTR flagged between 0.1% to 1.4% of spliced alignments in the assembled *A. thaliana* RNA-seq data as erroneous (**Supplemental Table S1.2**). The vast majority of these alignments (94.4% from HISAT2 and 96.9% from STAR) contained junctions not supported by the reference annotation. The remaining alignments contained junctions that were found in the annotation (49 and 117, for HISAT2 and STAR, respectively). The proportion of erroneous alignments varied across samples, with the *mddcc* mutant having over a fourfold increase in the proportion of erroneous alignments compared to the wild-type strain. In the *mddcc* mutant strain, EASTR flagged 138,999/32,614,412 (0.4%) erroneous alignments in HISAT2 and 257,794/18,652,175 (1.4%) in STAR, compared to 0.1% (31,249/33,230,121) and 0.2% (38,682/15,488,709) for the wild-type data. EASTR flagged a higher number of non-reference junctions in the *mddcc* alignments (5,734 in HISAT2 and 8,410 in STAR) than in the wild-type alignments (1,535 in HISAT2 and 1,425 in STAR). These observations were consistently observed across all samples (**Figure 2, Supplemental Table S1.2**).

#### Improving Transcript Assembly Quality

Just as with human and *Zea mays*, the application of EASTR filtering to alignments in the *A. thaliana* dataset improved the quality of transcriptome assembly, as shown in **Table 1**. We assembled transcripts with StringTie2 using both unfiltered and EASTR-filtered HISAT2 and STAR alignments, and compared the assemblies to the TAIR10.1 reference annotation. We observed a reduction in the number of non-annotated introns, exons, and transcripts per sample (**Supplemental Table S2.2**). The use of EASTR had a minimal impact on the number of reference-matching transcripts per assembly, with a marginal loss of ≤20 out of >15,000 total reference-matching transcripts per assembly for both HISAT2 and STAR (**Supplemental Table S2.2**). These observations were consistent in wild-type and mutant datasets.

#### Evaluating reference annotation

By applying EASTR to the TAIR10.1 annotation [23], we identified 283 introns within 316 transcripts and 193 genes that appeared to be potentially spurious (**Supplementary Table S4.6**). Consistent with our observations in *Z. mays*, we also identified instances of splicing between putative tandem gene duplications (**Supplemental Figure S4**) and potentially unannotated TEs. Our analysis also uncovered numerous annotation errors in several repeat-rich gene families, such as the Receptor-like proteins (RLP) with Leucine-rich repeat (LRR) domains [24]. This family encompasses 57 members, and we identified numerous spurious introns within this gene family (RLP18, RLP34, and RLP49). Within the polyubiquitin family, which contains tandem repeats of 228 bp encoding a ubiquitin monomer [25], we identified annotation errors in UBQ4, UBQ10, UBQ11, and UBQ14.

## Discussion

EASTR is a new computational tool that effectively identifies incorrect spliced alignments caused by repeat elements in RNA-seq datasets. By utilizing sequence similarity between the downstream and upstream sequences flanking a given splice junction, EASTR can identify and remove spuriously spliced alignments and also highlight potential errors in genome annotation, thereby improving the accuracy of downstream analyses that rely on alignment and annotation data.

In this study, we analyzed RNA-seq data from three species representing different tissue types and library preparation methods. Our analysis revealed that spurious alignments can account for up to 20% of the spliced alignments in the datasets we examined. The ribo-minus library preparation method had a higher proportion of spurious junctions and alignments compared to poly(A) selection, possibly because it captures nascent transcripts containing intronic sequences, which are typically enriched for repeat elements. In some samples, as many as 99.97% of the alignments that EASTR flagged for removal were not found in the reference annotation, suggesting they were likely spurious. Additionally, we observed a stark contrast in spurious alignments of reads sequenced from germline and somatic tissues of *Z. mays*, likely due to the different levels of TE expression in these two tissue types. Our findings underscore the importance of considering library preparation methods and tissue types when interpreting spliced alignment results, and demonstrate EASTR’s broad applicability in improving the accuracy of RNA-seq data alignment.

Our experiments also show that pre-filtering RNA-seq alignment files with EASTR can improve the accuracy of transcriptome assembly. We observed an increase in transcript assembly precision and a reduction in the count of novel (non-annotated) exons, introns, and transcripts in all samples.

Our EASTR-based analysis of reference gene catalogs illustrate how past errors in spliced alignment might have produced erroneous annotation that remains in these databases today. In all gene catalogs we examined, we found hundreds of likely cases of mis-annotation. One notable finding involved a transcript in the high-quality MANE human gene catalog containing an intron flanked by two Alu elements, an unlikely event requiring two consecutive exonization events (Supplemental Materials, Section 1). Accurate transcript annotation remains a challenge across all eukaryotic species, and the errors we observed here are likely to be repeated in many other genome annotation databases, in which EASTR has the potential to identify similar problems.

In conclusion, EASTR offers an effective solution for detecting spurious spliced alignments and annotation errors, and can substantially improve the accuracy of RNA-seq data alignment, transcript assembly, and annotation across diverse organisms and sequencing datasets.

Nonetheless, achieving precise transcript annotation remains challenging, particularly in species characterized by active transposons, high genomic TE composition, and frequent tandem gene duplication, underscoring the need for continued development of tools and methods to tackle this challenge.

## Methods

Splice-aware aligners may incorrectly map a read originating from a contiguous repeat element as a spliced alignment, especially if there are mismatches between the read and the reference genome. For example, if a read originating from a repeat sequence contains a mismatch to the reference genome, the aligner may attempt to optimize the alignment score by splicing it to a nearby, similar, repeat sequence **(Figure 5)**. Mismatches between RNA-seq reads and the reference genome are relatively common and can occur for various reasons, including variation between the individual and the reference genome, RNA editing events, and sequencing errors. A spurious spliced alignment of a read that originates from a repeat sequence can manifest in two ways: (1) the first part of the read is correctly aligned to one copy of the repeat, and the second part is spliced to another similar repeat nearby **(Figure 5A)**; or (2) the read comes from a repeat in one locus but has multiple mismatches compared to the reference genome, and the aligner finds a higher-scoring spliced alignment between two similar repeats in a different locus (**Figure 5B**).

**Figure 5.**
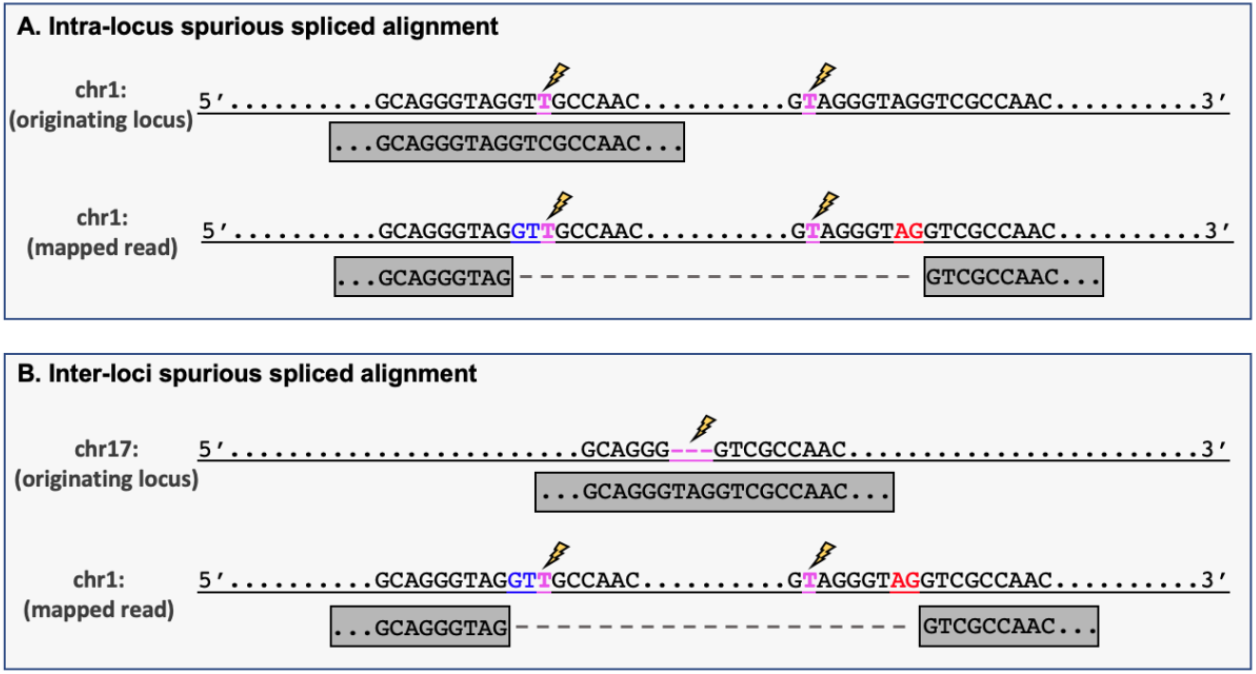
Schematic representation of spurious spliced alignment between repeat elements.**A. Intra-locus alignment error:** a read (gray box) originating from an upstream repeat element on chromosome 1 has a T to C mutation (highlighted in magenta and marked by a lightning bolt) relative to the reference genome. This read also has a single mismatch to the repeated downstream sequence. An aligner might erroneously create an intron between the two repeat elements, with the canonical GT-AG splice sites highlighted in blue and red. **B. Inter-loci alignment error:** a read (gray box) originating from chromosome 17 has a 3 bp insertion (highlighted in magenta and marked by a lightning bolt) relative to the reference genome. An aligner may align this read elsewhere in the genome and erroneously create an intron between two repeat elements.

### EASTR Algorithm for Detecting and Removing Spurious Spliced Alignments

EASTR aims to resolve the issue of erroneous splicing between repeat elements by recognizing sequence similarity between the flanking upstream and downstream regions of a specific intron, and the frequency of flanking sequence occurrence in the reference genome. The comprehensive workflow for detecting and removing spurious spliced alignments is described below and in **Figure 6**.

#### 1. Identification of potential repeat-induced spliced alignments

The input to EASTR is a set of alignments produced by a spliced aligner such as HISAT2 or STAR. For every intron identified in the input file, EASTR computes an alignment to identify similarity between an upstream “reference” sequence, centered on the 5’ splice site, and a downstream “query” sequence, centered on the 3’ splice site (**Figure 6**). The `mappy` Python wrapper of minimap2 [26] is used for this purpose, with k-mer length, minimizer window size, and chaining scores set to 3, 2, and 25, respectively. By default, EASTR extracts 100bp from both ends of the splice junction (SJ), extending 50bp in either direction from the splice site, to use as reference (upstream) and query (downstream) sequences. Following the alignment process, EASTR selects the primary `mappy` alignment if more than one good alignment was detected. The alignment is scored using a matrix that assigns 3 points for a match, 4 points penalty for a mismatch, 12 points penalty for opening a short gap, 32 points penalty for opening a long gap, 2 points penalty for extending a short gap, and 1 point penalty for extending a long gap. This scoring matrix is designed to be permissive in order to capture diverged homologous sequences, such as TE families, that may have sufficient homology to be erroneously splice-aligned due to stretches of exact matching bases.

**Figure 6.**
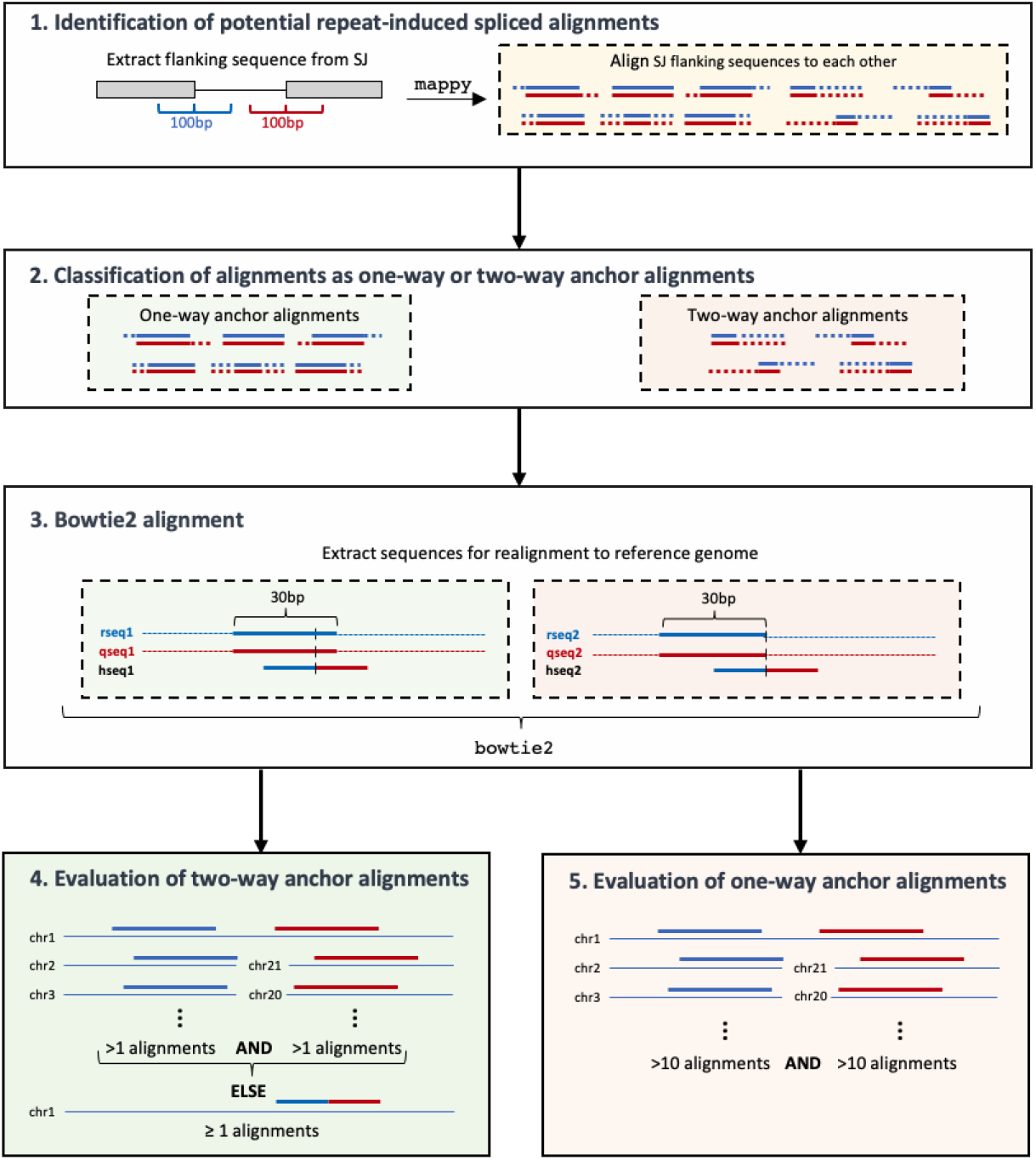
EASTR algorithm for detecting and removing spurious spliced alignments. The five-step process employed by EASTR to identify and filter erroneous splicing events caused by repetitive elements in a genome. (1) Identification of potential repeat-induced spliced alignments through preliminary alignments using mappy. Mappy is provided 100bp sequences centered on each end of the splice junction (SJ). (2) Classification of alignments as one-way or two-way anchor alignments based on the alignment of anchor sequences at a splice junction (3) Bowtie2 alignment to evaluate the occurrence frequency of reference, query, and hybrid sequences in the genome. (4) Assessment of two-way anchor alignments to determine the uniqueness of aligned sequences and classify spliced alignments as spurious or non-spurious. (5) Evaluation of one-way anchor alignments, considering duplicated exons, to identify and remove spurious junctions.

#### 2. Classification of alignments as two-way or one-way anchor alignments

EASTR examines `mappy` alignments generated in step 1, classifying each of them as either a “two-way anchor alignment” or a “one-way anchor alignment”. Anchors are short stretches of sequence that must align at the ends of a splice junction for the spliced alignment to be considered by the aligner. STAR and HISAT2 typically use minimum anchor sizes of 5-7bp for unannotated junctions. In EASTR, the default minimum anchor size is set to 7bp.

Two-way anchor alignments in EASTR meet the following criteria: 1) both reference and query alignment starting positions are less than or equal to the exon overhang (default: 50bp) minus the anchor length, or 43bp by default, 2) both reference and query alignment ending positions exceed the overhang plus anchor length (57bp by default), and 3) the alignment shift between reference and query sequences (absolute value of reference start minus query start) is below twice the anchor length (14bp by default; as illustrated in Figure 7A). Alignments not meeting these criteria are designated as one-way anchor alignments (Figure 7B).

**Figure 7.**
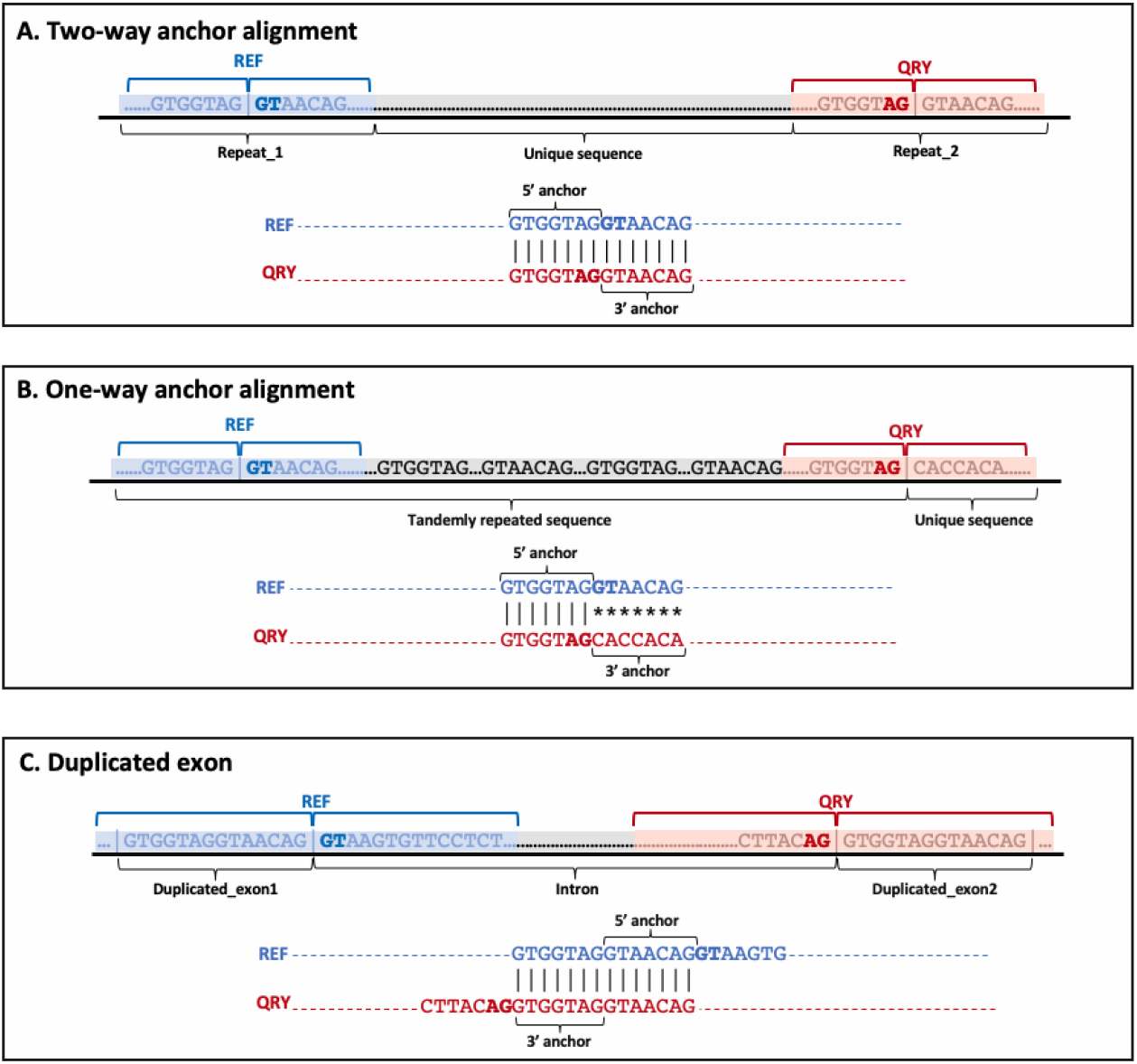
Examples of various alignments in the EASTR workflow. **(A)** Two-way anchor alignment with similar flanking sequences; **(B)** One-way anchor alignment with skewed similarity towards the 5’ end; **(C)** Duplicated exon scenario where alignment between the query and the reference sequences is primarily confined to exonic regions flanking the splice sites. The figure highlights the different scenarios that EASTR evaluates to detect and remove spurious spliced alignments caused by sequence similarity between upstream and downstream flanking regions.

#### 3. Bowtie2 alignment to check occurrence frequency

Following the initial `mappy` alignment, EASTR uses bowtie2 [25] to map the upstream “reference”, downstream “query”, and “hybrid” sequences (described below) back to the reference genome. This step is essential for detecting repetitive sequences whose high occurrence increases the likelihood of them causing erroneously spliced alignments. To perform this alignment, we extract three 30bp sequences: two from the center of the `mappy` alignment for the upstream and the downstream sequences and one hybrid sequence obtained by concatenating the 15bp upstream of the 5’ splice site with the 15bp downstream of 3’ splice site. Using bowtie2 with parameters “`-k 10 -- end-to-end -D 20 -R 5 -L 20 -N 1 -i S,1,0.50`” we map all three sequences back to the reference genome and count the number of times each aligns.

#### 4. Evaluation of two-way anchor alignments

For sequence pairs classified as two-way anchor alignments, EASTR uses `bowtie2` to further assess the uniqueness of the reference, query and hybrid sequences as described in step 3. If either the upstream or downstream alignment is unique and the hybrid sequence does not align elsewhere, the spliced alignment is deemed non-spurious. Conversely, if `mappy` finds an alignment and bowtie2 [27] aligns both sequences to more than one genomic location, EASTR marks the SJ as spurious.

#### 5. Evaluation of one-way anchor alignments

One-way anchor alignment between the reference and query flanking sequences could suggest a spurious spliced alignment. For instance, such cases can occur when there is sequence similarity between the 5’ ends of the reference and query sequences, but not between the 3’ ends (i.e., the repeat element causing the issue is not centered on the splice junction and is skewed toward the 5’ end on both ends of the splice junction, as illustrated in **Figure 7B**). In such situations, EASTR employs a two-step approach to address these scenarios:

##### a. Identifying duplicated exons

EASTR initially determines whether an alignment corresponds to a duplicated exon, which is not considered a spurious spliced alignment. In these cases, the alignment between the query and the reference sequences is primarily confined to exonic regions flanking the splice sites, leading to a shifted alignment of the query and reference sequences (**Figure 7C**). EASTR examines whether the query start site is shifted by ≥43bp (overhang of 50bp minus the anchor of 7bp). If such a shift occurs and the hybrid sequence is not present elsewhere in the genome, the alignment likely represents a pair of duplicated exons rather than a spurious junction.

##### b. Identifying spurious one-way anchor alignments

If the alignment does not meet the criteria for a duplicated exon, EASTR examines whether the upstream and downstream partially aligned sequences appear more than 10 times in the reference genome. If this is the case, the partial alignment is deemed spurious.

### Reference genomes and annotations

Human reads were aligned to the GRCh38 genome assembly after excluding pseudoautosomal regions and alternative scaffolds. The accuracy of the human transcriptome assemblies generated using StringTie2, as well as the non-reference and reference junction counts, were evaluated by comparing them to the GRCh38.p8 release of the RefSeq annotation, filtered to include only full-length protein-coding and long non-coding RNA transcripts. *A. thalian*a reads were aligned to TAIR10.1 (RefSeq accession GCF_000001735.4), and the accuracy of the transcriptome assemblies and junction counts were assessed by comparing them to the corresponding annotation [23]. *Z. mays* reads were aligned to the B73 NAM 5.0 assembly (RefSeq accession GCF_902167145.1) and the accuracies of the transcriptome assembly and junction counts were evaluated by comparing them to the corresponding NAM 5.0 Zm00001eb.1 annotation obtained from MaizeGDB [22]. The transposon annotation for *Z. Mays* was also retrieved from MaizeDGB.

### Alignment and assembly

A HISAT2 index was built using the following command: `hisat2-build -p 16 --exon genome.exon --ss genome.ss genome.fa hisat_index.` All RNA-seq datasets were aligned using HISAT2 [ref] with default parameters using the following command: `hisat2 -x hisat_index -1 R1.fastq -2 R2.fastq -S aligned.sam.` For *Arabidopsis* and maize lower leaf datasets, we added the `–rna-strandedness RF` flag to indicate an fr-firststrand library. Sorting and converting the resulting SAM files to BAM format was done with samtools [ref].

A STAR index was built using: `STAR --runThreadN 12 --runMode genomeGenerate --genomeDir star_index --genomeFastaFiles genome.fa --sjdbOverhang [read_length-1] --sjdbGTFfile reference.gtf`. RNA-seq datasets were aligned and sorted by STAR using the following command: STAR --runThreadN 12 --genomeDir star_index --readFilesIn R1.fastq R2.fastq --outSAMstrandField intronMotif --twopassMode Basic --outSAMtype BAM SortedByCoordinate --limitBAMsortRAM 16000000000 --outSAMunmapped Within --outFileNamePrefix sampleID [ref].

Transcriptome assembly was performed using StringTie2[ref] version 2.2.1, utilizing HISAT2 and STAR alignments, with the following command: `stringtie2 aligned.bam -o sample.gtf.`

### Assembly accuracy metrics

Sensitivity was quantified as the ratio of true positives (TP) to the sum of true positive and false negative (FN) exons, introns, or transcripts that match the reference annotation. Precision was quantified as the ratio of TP to the sum of TP and false positives (FP). True positives were defined as exons, introns, transcripts, or loci that match the reference annotation, and false negatives as exons, introns, transcripts, or loci in the annotation that were missing from the assembly. We used `gffcompare` [28] to count TP, FN, and FP as well as to profile the sensitivity and precision at the exon, intron, and transcript levels.

To assess the impact of EASTR on the number of novel and reference-matching introns, exons, and transcripts in each StringTie2 assembly, we calculated the percent change using the formula:

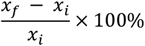

where x_i_ represents the count of introns, exons, or transcripts before applying EASTR to alignment files and x_f_ represents the count after assembly using EASTR filtered alignments. We employed the same percent change metric to evaluate the relative changes in transcript-level precision and sensitivity.

## Author Contributions

I.S. conceived the study, designed and implemented the software, conducted computational analyses, analyzed and interpreted the results, and wrote the manuscript. R.H. analyzed the results. H.J.J. and K-H.C. aided in the development of the software. M.P. conceived the study, advised the design and implementation of the software, and contributed to writing and editing the manuscript. All authors reviewed and approved the final manuscript.

## Acknowledgement

We thank Steven Salzberg for his helpful feedback and insightful comments on this manuscript. This work was supported in part by NSF grant DBI-1759518 and NIH grant R01-MH123567.

## Data Availability

The human DLPFC dataset is available from the NCBI Sequence Read Archive (BioProject; https://www.ncbi.nlm.nih.gov/bioproject/) under accession number PRJNA595606. The maize leaf dataset is available under accessions SRR10095075, SRR10095076, SRR10095077. The maize pollen dataset is available under accession numbers SRR3091548, SRR3091717, SRR3094513. The *Arabidopsis* dataset is available under accessions SRR14056780, SRR14056781, SRR14056782, SRR16596898, SRR16596890, SRR16596900.

## Code Availability

EASTR code and the analyses used in this paper are available through GitHub at https://github/ishinder/EASTR and https://github/ishinder/EASTR_analyses, respectively.

