## Supplemental Material for "EASTR: Correcting systematic alignment errors in multi-exon genes"

### Section 1: Questionable intron in TCEANC transcript

While the majority of MANE-selected isoforms serve as the preferred representatives for their corresponding genes, Sommer et al. identified several noteworthy exceptions [S1]. Our analysis proposes that TCEANC may be another such exception. Exonization events are infrequent, and consecutive Alu element exonization requires a minimum of four specific mutations [S2]. Moreover, Alu elements are susceptible to non-allelic homologous recombination (NAHR) [S3], which can produce deletions masquerading as introns during spliced alignments (**Figure S1C**), necessitating meticulous evaluation of intron splicing events. The GTEx RNA-seq data contain uniquely aligned spliced reads at this junction, possibly indicating a deletion. Although prevalent structural variant databases, such as dbVar [S4] and gnomAD [S5], report a chimeric Alu-producing duplication in this region (**Figure S1B**), further investigation is required to validate this junction, as additional recombinations, including deletions, may remain undetected due to factors such as structural variant caller limitations with short reads and the abundance of Alu elements. Furthermore, we utilized SpliceAI (version 1.3.1) to evaluate the acceptor and donor splice sites [S6]. The evaluation included the entire intron and an additional 200bp sequence upstream the donor and 200bp sequence downstream the acceptor. Following the guidelines from the SpliceAI manual (https://github.com/Illumina/SpliceAI), we included an extra 5,000bp on both the donor and acceptor side, resulting in a 10,000bp of flanking sequence context. Consequently, the full length of a splice site input into the SpliceAI model is 10,400bp plus the intron length. We calculated the average score from the five trained models for each site. The results, presented in **Figure S2**, demonstrate that the second putative exonization event is notably weak.

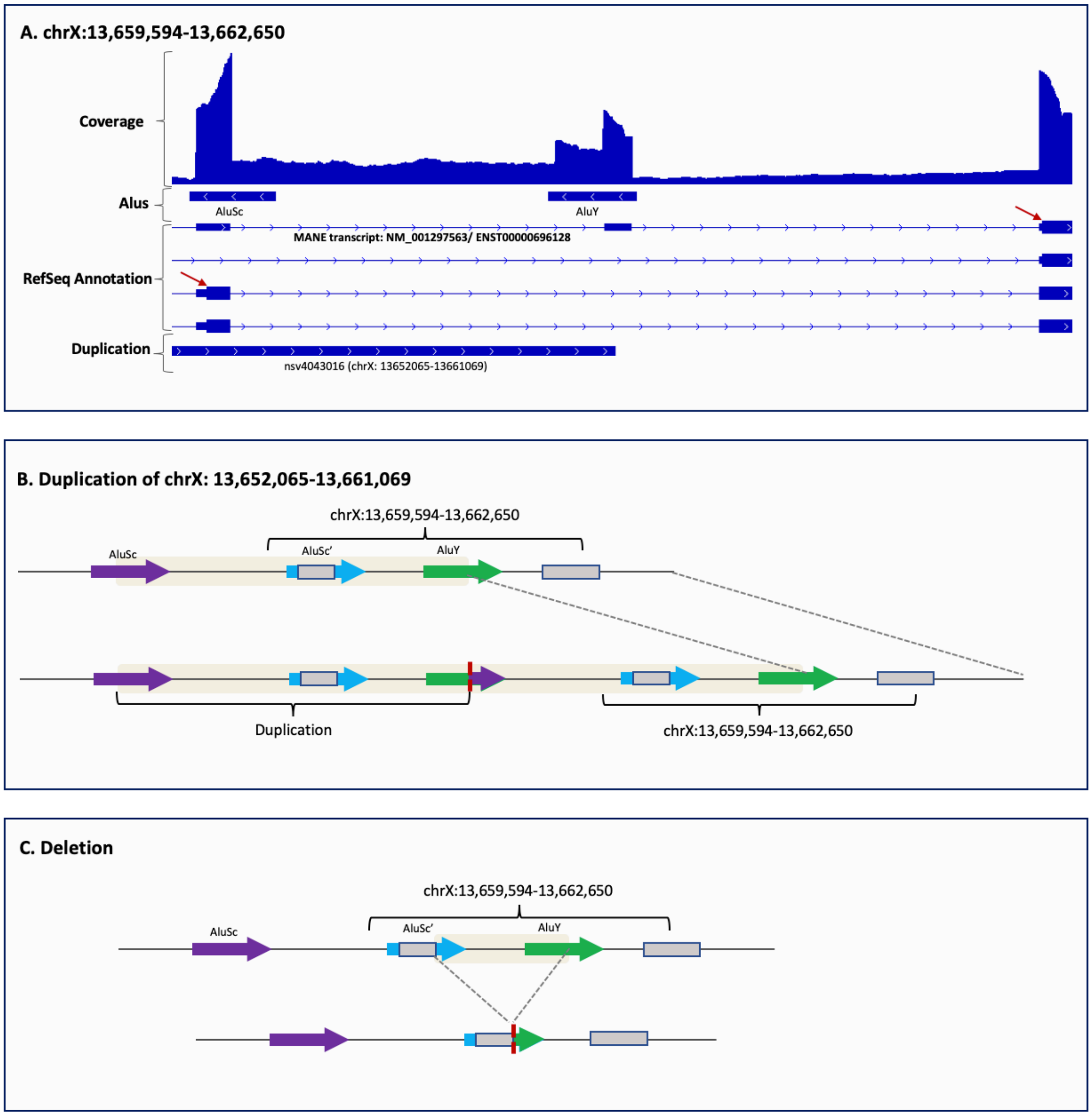


***Figure S1.*** Intronic Splicing Between Consecutive Alu Elements in TCEANC Gene: **(A)** The coverage track displays coverage from HISAT2 alignments across 9,795 GTEx samples [S7]. The *Alus* track illustrates two consecutive *Alu* elements, with the RefSeq annotation below. The MANE transcript includes exons situated within the consecutive *Alu* elements. Red arrows mark the start of an ORF. This figure emphasizes our analysis of the MANE catalog, where we identified a TCEANC gene transcript, specifically NM_001297563 (CHESS: CHS.57562.1, GENCODE: ENST00000696128), harboring an intron warranting further investigation. GTEx RNA-seq data reveals uniquely aligned spliced reads at this junction, potentially indicating a deletion. Common structural variant databases, such as dbVar and gnomAD, report a chimeric Alu-producing duplication in this region, as shown in the duplication track. **(B)** The impact of the duplication shown in (A) is schematically represented, displaying the formation of a chimeric Alu between an AluY (green arrow) in the TCEANC gene and an upstream AluSc (purple arrow). The blue arrow depicts an AluSc' element within the TCEANC gene. Arrows depict Alu elements, the yellow rectangle highlights the region of duplication, and the breakpoint is indicated by a red dashed line. Gray rectangles represent exons. **(C)** Schematic representation of a hypothesized deletion resulting from NAHR.

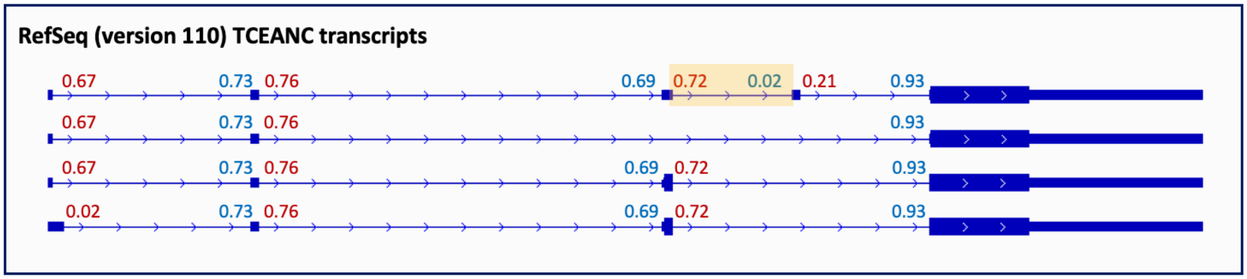


**Figure S2**. SpliceAI donor and acceptor scores for introns in the TCEANC gene. The figure displays all TCEANC RefSeq transcripts and the SpliceAI scores for donor (in red text) and acceptor (in blue text) splice sites. The questionable intron is highlighted in yellow. The SpliceAI acceptor site score for the questionable intron is 0.02, more than 30 times lower than all other acceptor scores in this gene. Moreover, the exon following the questionable intron exhibits a low donor site score. The combination of the low acceptor score at the 5' end and the low donor score at the 3' end of the exon supports our hypothesis that this splice site is likely spurious.

### Section 2: Incomplete Annotation and Erroneous Transcripts in the NPIPB3 Gene

The NPIP gene family, like the NBPF family examined in the results section, has expanded in primates through segmental duplications. The length of the VNTR region at the carboxy terminus is a defining characteristic that distinguishes paralogous copies [S8]. We found that none of the protein coding transcripts in RefSeq, CHESS, or GENCODE gene catalogs fully cover the characteristic VNTR region at the carboxy terminus of NPIPB3 (**Figure S3**). Furthermore, this protein-coding gene is entirely absent from the MANE gene catalog. The sawtooth pattern observed in the coverage of spliced alignments suggests the presence of many erroneous spliced alignments in this repetitive region. Our findings are supported by the UCSC genome browser visualization of the CHM13 liftOver track, which reveals a nearby 252bp deletion in CHM13 relative to GRCh38. This deletion exactly matches the length of the intron in RefSeq transcript NM_130464 (CHESS3 transcript CHS.19306.24), strongly suggesting that it is a VNTR indel.

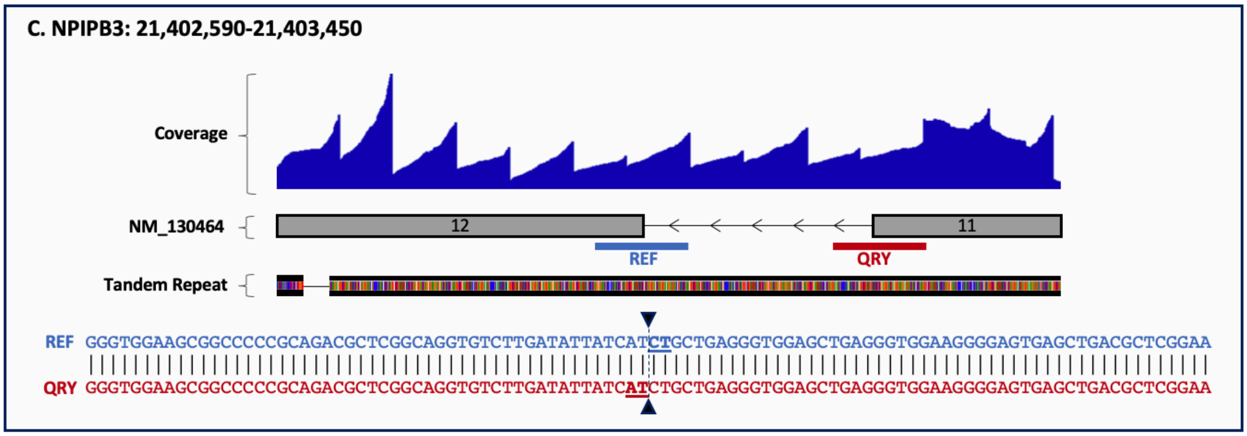


**Figure S3.** Error in annotation of NPIPB3 transcript NM_130464 caused by a VNTR polymorphism. A 252bp intron is incorrectly inserted between exons 11 and 12. The 100bp alignment presented below the transcripts displays identical upstream and downstream sequences flanking the intron in NM_130464, with the splice site indicated by inverted triangles. The erroneous intron's length is twice the size of the 126bp tandem repeat. The coverage track presents the coverage from 9,795 GTEx samples [S7].

### Section 3: Errors in TAIR 10.1 gene annotation

**Figure S4** presents an example of erroneous splicing between putative tandem gene duplications, highlighting the need for careful evaluation and refinement of gene annotation methods.

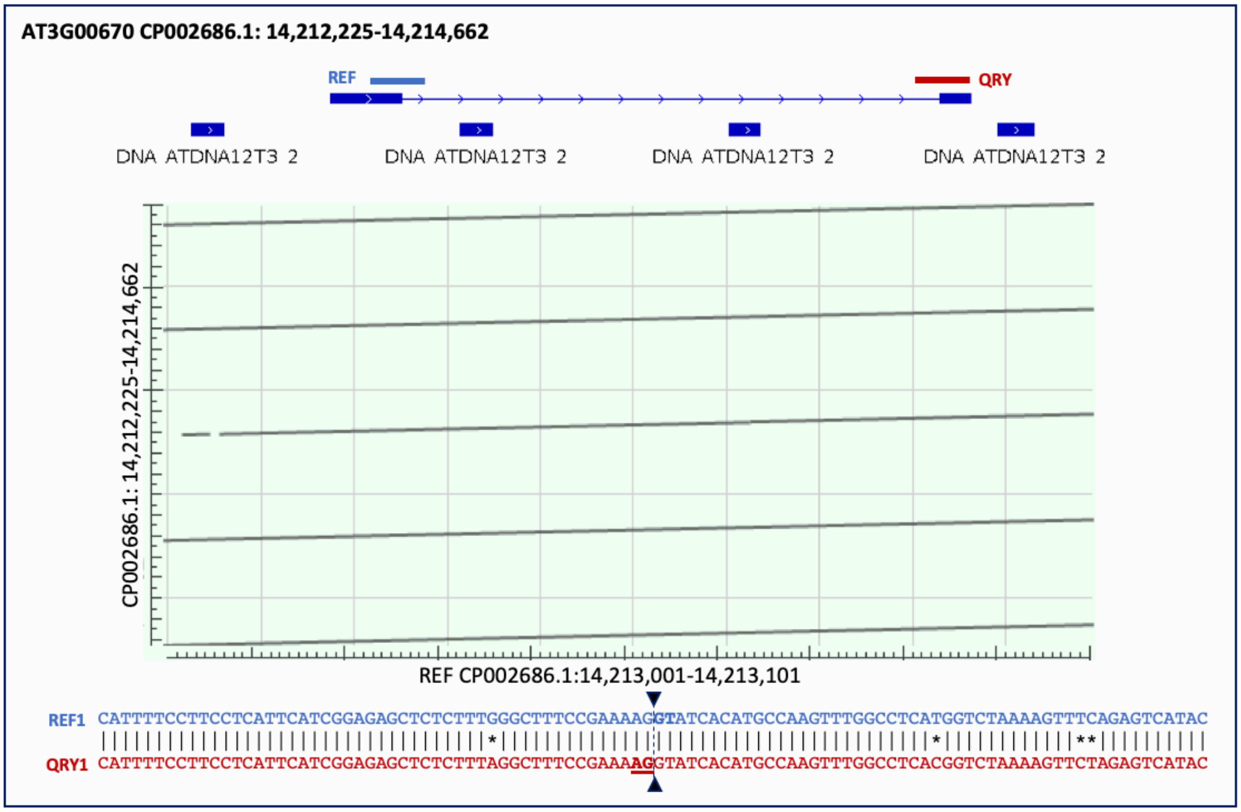


***Figure S4*. Erroneous Splicing between tandemly duplicated regions in A. thaliana:** The top track displays the reference transcript for gene AT3G00670. The TE annotation track beneath the transcript illustrates tandemly duplicated transposons. The dot plot, generated using NCBI blastn[S9] and located below the transcript and TE annotation tracks, features the reference query (REF) on the x-axis and the region presented in the transcript track on the y-axis. The reference sequence is observed to appear four times in this region. Alignments of the upstream and downstream intron-flanking sequences (REF and QRY) in transcripts exhibit 96% homology.
